# Goal-directed modulation of stretch reflex gains is reduced in the non-dominant upper limb

**DOI:** 10.1101/2022.11.30.518527

**Authors:** Frida Torell, Sae Franklin, David W. Franklin, Michael Dimitriou

## Abstract

Most individuals experience their dominant arm as being more dexterous than the non-dominant arm, but the neural mechanisms underlying this asymmetry in motor behaviour are unclear. Using a delayed reach task, we have recently demonstrated strong goal-directed tuning of stretch reflex gains in the dominant upper limb of human participants. Here, we used an equivalent experimental paradigm to address the neural mechanisms that underlie the preparation for reaching movements with the non-dominant upper limb. We found only minor goal-directed differences in the short latency stretch reflex of the non-dominant limb. There were consistent effects of load, preparatory delay duration and target direction on the long latency stretch reflex. However, by comparing stretch reflex responses in the non-dominant arm with those previously documented in the dominant arm, we demonstrate that goal-directed tuning of short and long latency stretch reflexes is markedly weaker in the non-dominant limb. The results indicate that the motor performance asymmetries across the two upper limbs is partly due to the more sophisticated control of reflexive stiffness in the dominant limb, likely facilitated by the superior goal-directed control of muscle spindle receptors. Our findings therefore suggest that independent fusimotor control plays a role in determining performance of complex motor behaviours and support existing proposals that the dominant arm is better supplied for executing more complex tasks, such as trajectory control.

**Key points:**

- Most of us routinely rely on the dominant arm to perform more complex and demanding motor tasks, but the mechanisms enabling the superior motor performance of the dominant limb are unclear.
- A better understanding of the motor asymmetry across the two arms might provide key insight into core sensorimotor principles.
- This study shows that goal-directed tuning of short and long latency stretch reflexes in the non-dominant arm is markedly weaker than in the dominant arm.
- Our results suggest that the more sophisticated control of reflexive stiffness in the dominant limb, likely facilitated by superior fusimotor control, partly underpins the laterality of motor performance.

## Introduction

Most people display significant arm dominance, often termed ‘handedness’, which can be expressed as superior reaction time, dexterity, and strength in the dominant arm (Annett *et al.*, 1979; Goble & Brown, 2008a; Shen & Franz, 2005). Hand movement performance variations have been related to differences in the neurological organization of the sensorimotor system. It was initially suggested that the observed asymmetries in motor behaviour are related to the cortical hemisphere controlling movement of the arm (Liepmann, 1908). Later studies have demonstrated that both hemispheres contribute significantly to the control of contralateral goal-directed movements (Fisk & Goodale, 1988; Haaland & Harrington, 1989; Winstein & Pohl, 1995), although it has more recently been proposed that the left-hemisphere in right-handers plays a specialized role in the control of complex ipsilateral and contralateral movements (Haaland & Harrington, 1996).

The hand dominance displayed by humans has led to a more specific theory of lateralization of motor control processes (Sainburg, 2005; Sainburg, 2014; Sainburg & Kalakanis, 2000). This proposal suggests that the dominant arm is primarily controlled through feedforward control of trajectory with accurate internal models of limb dynamics, whereas the non-dominant arm is primarily controlled through impedance control for stabilization. Such a theory can nicely explain why we might choose specific hands for certain tasks during bimanual object manipulation. For example, when unscrewing the lid of a jar, people will often hold (stabilize) a jar with their non-dominant hand and use the dominant hand to open the lid. Similarly, when threading a needle, the non-dominant hand might stabilize the needle while the dominant hand provides the fine manipulation of the thread. Indeed, it has been shown that, in certain contexts, the non-dominant arm exhibits more effective load compensation responses than the dominant limb (Bagesteiro & Sainburg, 2003). It has been suggested that such limb dominance might reflect differences in the hemispheric control of the body, where each hemisphere is specialized for different but complementary functions: the dominant system for controlling limb trajectory dynamics, and the nondominant system for controlling limb position (Sainburg, 2005).

Little attention has been directed to whether differences in dexterity across the arms can partly represent differences in the goal-directed control of stretch reflexes. Recent evidence has shown that muscle spindle firing is highly task- or goal-dependent, e.g., (Dimitriou, 2016; Papaioannou & Dimitriou, 2021; Ribot-Ciscar *et al.*, 2009). However, postural studies of reflex modulation have found little or no lateralization or specialization of stretch reflex responses between the two limbs (Maurus *et al.*, 2021; Walker & Perreault, 2015). Nevertheless, there is evidence that posture and movement exhibit different control strategies (Kurtzer *et al.*, 2005; Scheidt & Ghez, 2007). Moreover, it has been shown that both proprioceptive stretch reflexes and visuomotor feedback responses are selected according to the upcoming movement (Cesonis & Franklin, 2022; Ahmadi-Pajouh *et al.*, 2012; Wagner & Smith, 2008; De Comite *et al.*, 2021; Maeda *et al.*, 2021).

It has long been shown that long latency stretch reflexes exhibit task-dependent and goal-directed modulation (Hammond 1956; Crago *et al.*, 1976; Akazawa *et al.*, 1983; Kimura *et al.*, 2006; Pruszynski *et al.*, 2008; Nashed *et al.*, 2014). However, our recent work has shown that sufficient preparation time allows the goal-directed tuning of both short and long latency reflex responses of the dominant upper limb (Papaioannou & Dimitriou, 2021, Torell *et al.*, 2023). Here, we hypothesize that lateralization of independent fusimotor control might affect the goal-directed modulation of stretch reflexes in the non-dominant limb. That is, we predict that some of the behavioural differences resulting from limb dominance may reflect differences in the control of the gamma motor neuron system, which would result in differences in the goal-directed modulation of stretch reflexes. We test this hypothesis by examining the preparatory goal-directed tuning of stretch reflex gains in the non-dominant upper limb. We use a delayed-reach (center-out) task to examine the stretch reflex responses of the non-dominant arm under different background loads and preparatory delays. The current study was specifically designed to reveal whether the non-dominant arm displayed similar benefits of assistive loading and sufficient preparation time as means to promote the goal-directed tuning of stretch reflex gains, as shown previously to be the case for the dominant limb (Torell *et al.*, 2023).

## Materials and Method

### Participants

All participants involved in this study were right-handed and neurologically healthy. We recorded muscle activity from the non-dominant left arm of 16 individuals (mean age 26.8 years, SD 6.4 years; 8 were female). All participants gave informed, written consent prior to participating in the study, per the Declaration of Helsinki. This experiment was part of a research program approved by the Ethics Committee of Umeå University, Umeå, Sweden. All participants were financially compensated for their contribution to this study. The current study also contrasted equivalent muscle responses previously recorded from the right dominant arm of 14 additional individuals (Torell *et al.*, 2023). Although using two groups of right-handed participants to assess the impact of handedness on reflex tuning could be conceived as potentially limiting in terms of statistical power, we have no reason to believe that our two random population samples differed in an important way in terms of their ‘right-handedness’.

### Robotic platform

To perform the recordings from the left non-dominant arm, each participant sat upright in a customized chair that was mechanically stabilized to the floor. The chair was placed in front of a Kinarm™ robotic platform (Kinarm end-point robot, BKIN Technologies Ltd., Canada). The participants used their left hand to grasp the left handle of a bimanual robotic manipulandum (Fig. 1). Their forearm was placed inside a customized foam-cushioned structure made of hard plastic material and secured in this structure using a leather fabric with Velcro attachments. The attachment ensured a tight mechanical connection between the participant’s forearm, the plastic structure and the robotic handle. Velcro attachments were also used to secure an airsled to the plastic structure enveloping the forearm. The airsled allowed frictionless movement of the arm in a 2D plane. The robotic platform recorded kinematic data and sensors inside the handle (six-axis force transducer; Mini40-R, ATI Industrial Automation, U.S.A.) recorded forces exerted by the participants’ hand. Kinematic and force data from the Kinarm platform were sampled at 1 kHz. The same setup with the right handle was previously used for generating equivalent data from the dominant right arm of a separate group of participants (Torell *et al.*, 2023).

**Fig. 1.**
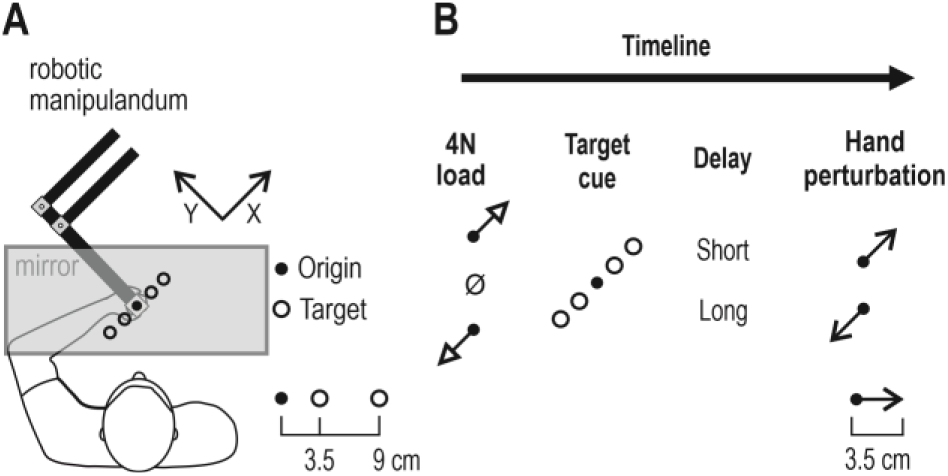
The robotic platform and experimental setup. **(A).** The participants were seated in an adjustable chair placed in front of the robotic platform. The participant’s non-dominant left hand grasped a robotic handle. All visual stimuli were projected onto a one-way mirror. The participant could not view their hand or the robotic handle in this setup. The position of the hand was represented by a white cursor. Four visual targets were continuously displayed along the X-axis. Two ‘near’ and two ‘far’ targets were used, placed at 3.5 cm and 9 cm from the origin, respectively. **(B)** In this experiment, each trial was initiated when the participant brought the cursor to the origin. A slow-rising 4N load in either the +X (upper right direction) or -X direction (lower left direction) or a ‘null’ load (no load) was then applied. Regardless of load, the participant had to keep the hand at the origin. One of the four targets was then cued by turning red. The target remained in the cued state for either 250 ms or 750 ms. This preparatory delay was followed by a rapid position-controlled perturbation of the hand (3.5 cm in 150 ms) in either the -X or +X direction. The cursor position was frozen during the perturbation. At the end of the perturbation, the target turned green (‘Go’ signal), and the participant had to swiftly move the hand to the target.

### Experimental design

All visual stimuli were projected on a one-way mirror which otherwise occluded view of the hand and robotic handle (Fig. 1A). Four orange circle outlines (‘targets’; 2.4 cm diameter) and a circle outline representing the origin (1.3 cm diameter) were situated along the X-axis (as defined in Fig. 1A). The center-to-center distance between the origin and the two ‘near’ targets was 3.5 cm, and the distance between the origin and the two ‘far’ targets was 9 cm. Hand position was represented by a white dot (‘cursor’; 1 cm diameter). To start a new trial, the participant had to move the white cursor to the origin. There, they had to remain immobile for a random wait period of 1-1.5 seconds. Provided that they had remained immobile inside the origin, the robotic manipulandum then generated a slow-rising 4N load in either the +X or -X direction or no such pre-load was produced (Fig. 1B). Any slow-rising load had a rise-time was 800 ms and a hold-time was 1200 ms.

At the end of this hold-time, and if the cursor remained immobile inside the origin, one of the four targets was cued by turning into a filled red circle and the target remained in this cued state for a relatively short (250 ms) or long time period (750 ms). At the end of this preparatory delay, a position-controlled haptic perturbation was delivered in either the +X or -X direction (3.5 cm displacement, 150 ms rise-time, and no hold). That is, the hand was perturbed towards or away from the cued target. The cursor position was frozen for the duration of the perturbation. The haptic perturbations were designed to induce kinematics of a fast naturalistic point-to-point movement; that is, the resulting velocity profiles were approximately bell-shaped. To achieve the desired hand kinematics on each trial, the robotic platform was allowed to apply the appropriate stiffness with as maximum of ~40,000 N/m, regardless of load/force conditions. Once the mechanical perturbation ended (i.e., 150 ms after perturbation onset) the filled red circle (i.e., cued target) turned green, representing the ‘Go’ signal to reach the target. The participants were instructed to swiftly bring the cursor to the highlighted target.

After reaching the target, the participants were required to keep the hand at the target for 300 ms. Thereafter, the participants received visual feedback on the performance on this specific trial. The visual feedback was ‘Too Slow’, if the time between the onset of the ‘Go’ signal and the time the cursor entered the highlighted target was >1200 ms, ‘Correct’ if the target was reached after 400-1200 ms and ‘Too fast ‘ if the target was reached after <400 ms. The selected feedback intervals encouraged the participant to move swiftly to the target, while allowing enough time for reaching the far target even when the hand had been perturbed in the opposite direction. The feedback message was visible for 300 ms. The participant then returned the cursor to the origin to initiate the next trial. If the participant wanted a break they could move their hand to the side, instead of returning to the origin circle. Breaks were normally encouraged every two blocks of trials, where a ‘block’ represents one set of the 48 unique trials. Breaks normally lasted <5 min. The experiment contained 48 unique trial types: 4 targets (two directions and two distances) x 3 load conditions (slow-rising 4N in the +X direction, -X direction or null) x 2 delays (250 ms and 750 ms) x 2 perturbation directions (+X and -X). There were 15 repetitions of each trial type (i.e., total number of trials was 720), and the trials were presented in a block-randomized manner. Each experiment took approximately 1.5 hours to complete. The data recorded using this experimental design were analysed separately in the current study, but also contrasted with previously recorded data from the dominant right arm. The data from the dominant right arm were generated using a mirror-equivalent version of experiment described above (Torell *et al.*, 2023).

### Electromyography (EMG)

Surface EMG electrodes (Bagnoli™ DE-2.1, Delsys Inc., USA) were placed on seven arm muscles: (1) *m. brachioradialis,* (2) *m. biceps* brachii, (3) *m. triceps brachii caput laterale,* (4) *m. triceps brachii caput longum,* (5) *m. deltoideus pars anterior,* (6) *m. deltoideus pars posterior,* and (7) *m. pectoralis major.* Prior to electrode placement, the skin was cleaned using alcohol swabs and the electrodes were coated with conductive gel. The electrodes were secured using surgical tape. The ground electrode had a diameter of 5.08 cm (Dermatrode® HER Reference Electrode type 00200-3400; American Imex, Irvine, CA, USA), and was placed on the *processus spinosus* of C7 region. The EMG signals were band-pass filtered online through the EMG system (20 – 450 Hz) and sampled at 1.0 kHz.

### Data pre-processing

Data pre-processing was performed using MATLAB® (version R2020b, MathWorks, Natick, MA, USA). EMG data was high pass filtered using a fifth-order, zero phase-lag Butterworth filter with a 30 Hz cut-off and then rectified. Movement onset of each trial was defined as the time when 5% peak velocity was reached. The EMG data was normalized (z-transformed) to allow for comparisons of separate muscles and participants. The procedure to z-transform the data has been described in detail in elsewhere (Dimitriou, 2014, 2016, 2018). In short, the z-transformation is performed by concatenating all EMG data from one muscle (including activity during voluntary movement) and calculating a grand mean and grand standard deviation across all raw data. Each muscle’s grand mean is then subtracted from the measured EMG values which are then divided by the grand standard deviation. An additional EMG normalization approach was used to further evaluate reflex tuning differences between the dominant and non-dominant arm. This approach involved producing the average (mean) unnormalized EMG trace across repetitions of a trial type (aligned on perturbation onset) for each separate muscle/participant. For each muscle, the maximal EMG value across all averaged traces (i.e., across all trial types) observed anytime 50 ms after perturbation onset was used to normalize the raw EMG data as a proportion (%) of this value. As described in the Results section, equivalent statistical results were obtained regarding differences in reflex tuning between the dominant and non-dominant arm, regardless if the %-normalized or z-normalized approach was used.

Throughout, the first five blocks of trials were viewed as familiarization trials and were not included in the subsequent analyses. For each participant, data was averaged for each trial type for plotting and statistical analyses. For plotting purposes only, signals were smoothed using a 5 ms moving window. The data pre-processing procedures were identical to those previously used with regard the right dominant arm (Torell *et al.*, 2023). In addition to investigating the reflex modulation of the non-dominant arm, the current study also compared the goal-directed tuning of reflexes between the non-dominant (left) and dominant (right) upper limbs. Data from the dominant right arm were generated in a previous study, where a separate group of participants performed the mirror equivalent experiment to the current one using their right dominant arm (Torell *et al.*, 2023). Because target direction was the primary factor shaping reflex gains in both the dominant (and non-dominant arm), the data used for these specific analyses were collapsed across target distances, so that ‘goal’ only represented target direction. To produce a measure that is more directly representative of goal-dependent tuning, the responses observed when preparing to stretch the homonymous muscle (i.e., trials where reaching the target required lengthening of the muscle) were subtracted from the responses observed when preparing to shorten the homonymous muscle.

### Statistical analyses

Statistical analyses were performed using z-normalized EMG data. Since the scope of the study was to investigate stretch reflex responses, only EMG data from muscles that were stretched by the haptic perturbations were examined. Across participants, we generated median EMG values for three predetermined epochs: pre-perturbation epoch (25 ms period prior to perturbation onset), the SLR epoch (25-50 ms period post perturbation onset) and LLR epoch (75-100 ms post perturbation onset). Specifically, the analysed data contained the median EMG signal for each trial (i.e., median of each load, delay and target combination), for each muscle and participant in the three aforementioned epochs. To study the main and interaction effects of the non-dominant arm, the median EMG data was used to perform repeated-measures analysis of variance (ANOVA) of the design 2 (delay) x 3 (loads) x 2 (target distances) x 2 (target directions), separately for each muscle type. For the SLR response in particular, ANOVA was conducted without including the pre-loaded (‘loaded’) muscle condition as it is known that automatic gain-scaling accompanying pre-loading tends to saturate the SLR response, preventing its goal-directed modulation (as occurred in our study; e.g., Figures 3-4). Post-hoc analyses were performed using Tukey’s HSD (honest significant difference). To check normality Shapiro-Wilks test for samples with <50 data-points and Lilliefors test for larger samples were used. Statistical comparisons of reflex responses between dominant and non-dominant limb were conducted with independent t-tests. All statistical analyses were performed or STATISTICA® (StatSoft Inc, USA).

The onset of SLR reflex modulation was estimated using receiver-operator characteristic (ROC) technique (Green & Swets, 1966). The ROC area is value that assesses the overall performance of a binary classifier, where a ROC area of 1 and 0 represents perfect discrimination and a ROC area of 0.5 represents a discrimination performance equal to chance. Since the differences between the target directions were stronger for the longer preparatory delay, we used this data for the ROC analysis. Specifically, the EMG curves of targets in the direction of homonymous muscle stretch were contrasted to EMG curves of targets in the direction of muscle shortening. Averages across target distance were viewed as representative of the reflex modulation in the population. In accordance with the results of Corneil *et al.*, the discrimination was viewed as significant when the ROC area remained >0.75 for five consecutive time periods (i.e., for at least 5ms) (Corneil *et al.*, 2004). To assess individual reflex modulation onset, a similar approach was used on individual EMG responses across trials. This was done separately for each participant. To eliminate the risk of false positives, each SLR modulation onsets was confirmed by visual inspection. The ROC curves were created using MATLAB® (MathWorks R2020b, Natick, MA, USA).

## Results

### The non-dominant arm

Sixteen participants were recruited to perform a delayed-reach task using their non-dominant left arm. On each trial, one of three different loads was applied (−4N, 0N or 4N), and the participants had to maintain their hand positioned inside the origin regardless. One of four targets was then cued (turned red), and after either a short or long preparatory delay the left hand was perturbed by 3.5 cm in one of two directions (+X or -X; Fig. 1). At the end of the haptic perturbation, the colour of the cued target changed from red to green, indicating to participants that they need to actively reach the cued target. This experiment examines how the target location (direction and distance), background load and delay between the presentation of the target and the perturbation impact reflex responses during reach preparation.

Figure 2 illustrates responses from the pectoralis muscle of a single participant. Visual inspection of the figure points to clear differences in EMG as a function of target direction during the long-latency reflex epoch (LLR; 75-100 ms). However, there is no clear differentiation of EMG as a function of target direction during the short-latency reflex epoch (SLR; 25-50ms). Figures 3-4 represent equivalent averaged responses across all 16 participants, for the pectoralis and posterior deltoid, respectively. In contrast to what we have observed for the dominant right arm (Torell *et al.*, 2023), our current figures and analyses indicate that goal-directed tuning of SLR responses was not prevalent in the non-dominant left arm. Specifically, in what follows, we first examine the impact of target location, load and delay on reflex responses of the left arm and then compare the level of goal-directed tuning of reflexes in the left and right arm. The statistical analyses confirm our initial hypothesis of reduced goal-directed tuning of stretch reflexes in the non-dominant upper limb.

**Fig. 2.**
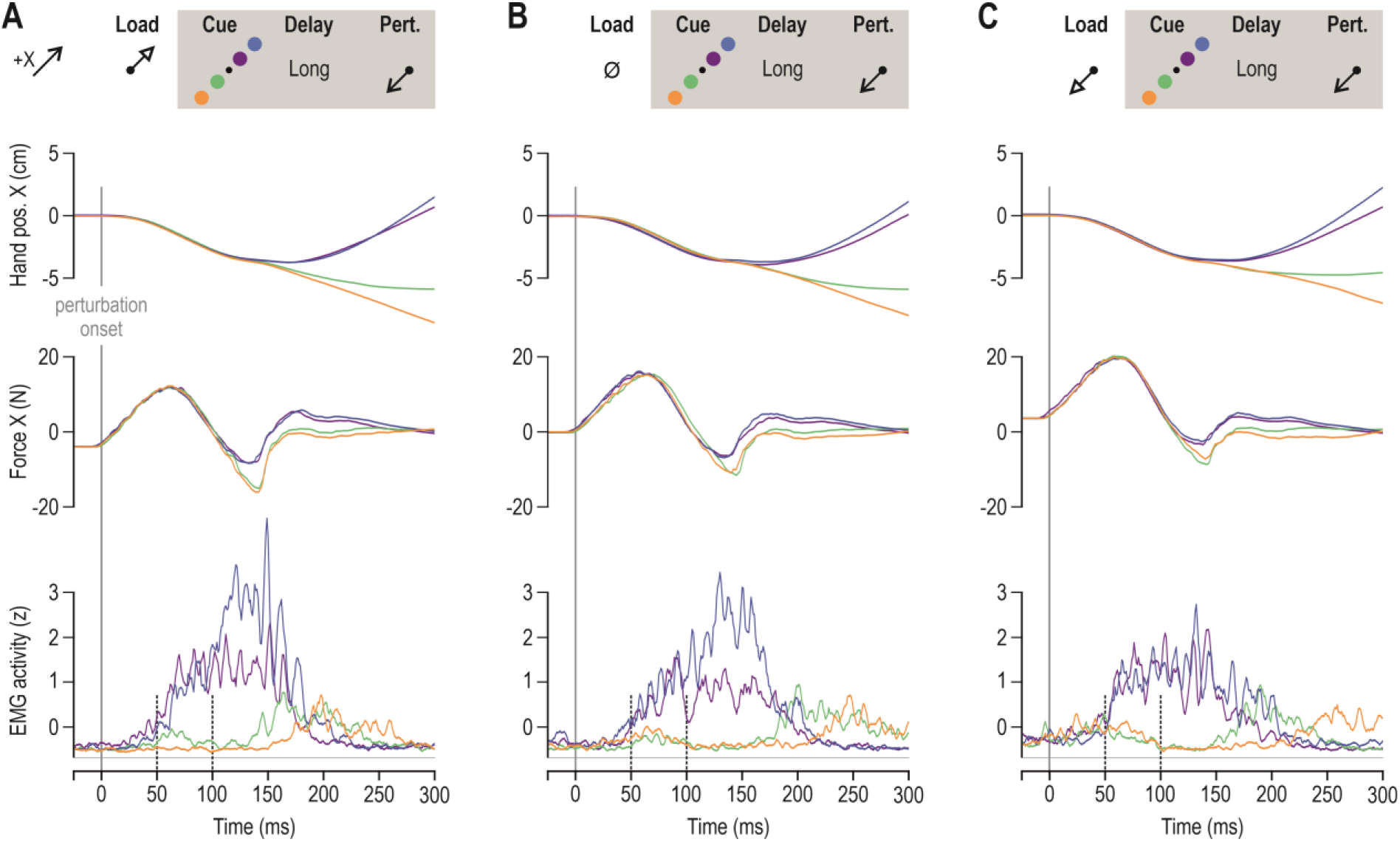
Responses from the pectoralis muscle of a single participant in the delayed reach task. Blue and purple traces represent trials where ‘far’ and ‘near’ targets were placed in the +X direction, respectively. These targets require shortening of the pectoralis in order to be reached. Orange and green traces represent trials where ‘far’ and ‘near’ targets were placed in the -X direction, respectively. These targets require stretch of the pectoralis in order to be reached. All data in this figure represent median responses across relevant trials that involved a long preparatory delay (750 ms). **(A)** A slow-rising load was first applied in the +X direction, unloading the pectoralis prior to the haptic perturbation, the onset of which is signified as time zero **(B)** As ‘A’, but no load was applied prior to the haptic perturbation **(C)** As ‘A’, but the slow-rising load was applied in -X direction, loading the pectoralis prior to stretch.

**Fig. 3.**
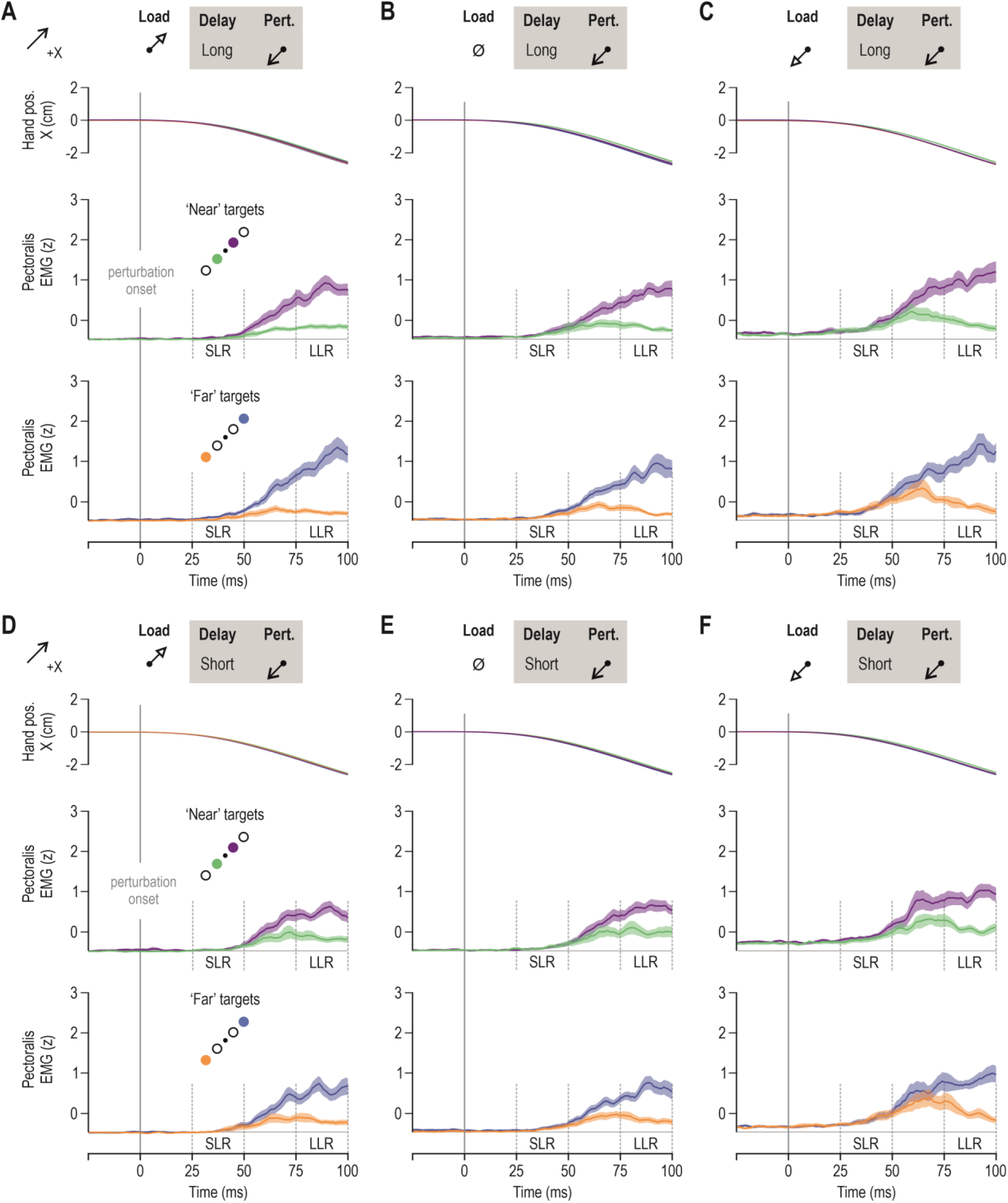
Stretch reflex responses in the pectoralis major of the non-dominant limb. Throughout, blue and purple traces represent trials where participants had to reach the ‘far^1^ and ‘near’ targets in the +X direction, respectively. Reaching these targets required shortening of the pectoralis. Orange and green traces represent trials where the participant had to reach the ‘far’ and ‘near’ targets in the -X direction, respectively. These trials require stretch of the pectoralis. Throughout, colour shading represents ±1 SEM. **(A)** The upper panel represents mean hand position across participants (N = 16), for all trials where the pectoralis muscle was unloaded before being stretched by the haptic perturbation, following a long preparatory delay (750 ms). The middle row displays mean pectoralis EMG activity across participants for the subset of trials where one or the other ‘near’ targets were cued for a long delay before pectoralis stretch; the bottom panel represents the equivalent for ‘far’ targets. **(B)** As ‘A’, but representing the ‘no-load’ trials **(C)** As ‘A’ but representing trials where the pectoralis was loaded before the stretch perturbation. **(D-F)** As ‘A-C’ but representing trials where the preparatory delay was relatively short (250 ms).

**Fig. 4.**
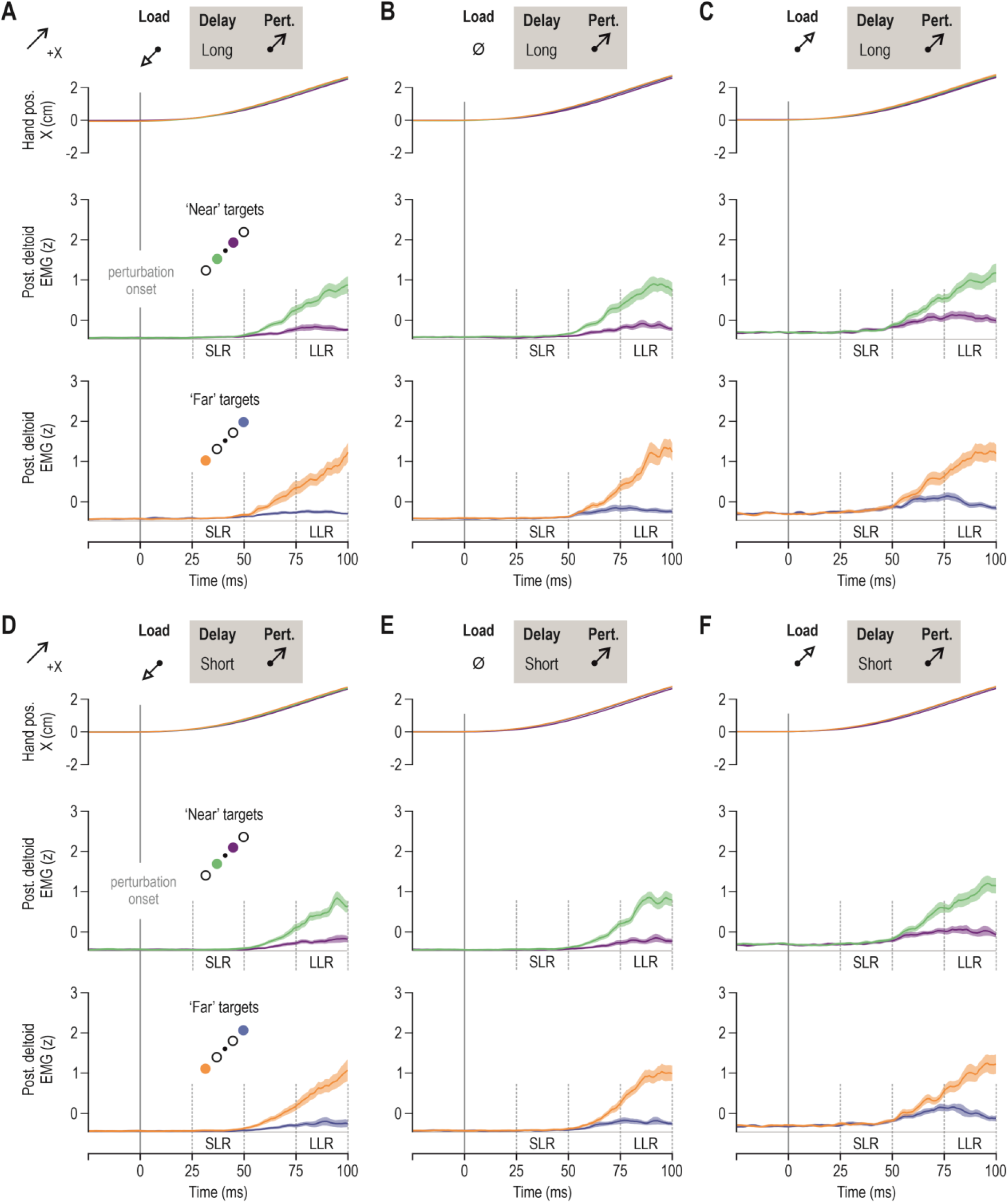
Stretch reflex responses in the posterior deltoid of the non-dominant limb. Throughout, blue and purple traces represent trials where participants had to reach the ‘far’ and ‘nea? targets in the +X direction, respectively. Reaching these targets required stretch of the posterior deltoid. Orange and green traces represent trials where the participant had to reach the ‘far’ and ‘near’ targets in the -X direction, respectively. These trials require shortening of the posterior deltoid. Throughout, colour shading represents ±1 SEM. **(A)** The upper panel represents mean hand position across participants (N = 16), for all trials where the posterior deltoid muscle was unloaded before being stretched by the haptic perturbation, following a long preparatory delay (750 ms). The middle row displays mean posterior deltoid EMG activity across participants for the subset of trials where one or the other ‘near’ targets were cued for a long delay before posterior deltoid stretch; the bottom panel represents the equivalent for ‘far’ targets. **(B)** As ‘A’, but representing the ‘no-load’ trials **(C)** As ‘A’ but representing trials where the posterior deltoid was loaded before the stretch perturbation. **(D-F)** As ‘A-C’ but representing trials where the preparatory delay was relatively short (250 ms).

#### The pre-perturbation epoch

The pre-perturbation epoch (i.e., 25 ms period before perturbation onset) was characterized by significant main effects of load in all three analysed muscles. Specifically, as expected, there was significantly higher pectoralis EMG activity when the muscle was loaded, with the lowest pre-perturbation values observed when the muscle was unloaded (F_2,30_ = 80.1, p <10^-5^, 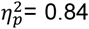; Tukey’s HSD loaded vs. no load: p = 0.0001; loaded vs. unloaded: p = 0.0001; no load vs. unloaded: p=0.0024). For the anterior deltoid, there was also a main effect of load on pre-perturbation EMG activity (F_2,30_ = 6.9, p = 0.0034, 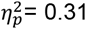) with Tukey’s HSD test indicating significantly higher values when the muscle was loaded (vs. unloaded: p=0.01; vs. no load: p=0.007). Finally, the posterior deltoid also showed a main effect of load on pre-perturbation activity (F_2,30_ = 36.3, p <10^-5^, 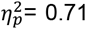) with Tukey’s HSD test indicating significantly higher values when the muscle was loaded (vs. unloaded: p=0.0001; vs. no load: p=0.0001).

For the pectoralis muscle alone, there was also a main effect of target direction on the pre-perturbation EMG activity (F_1,15_ = 10.2, p = 0.006, 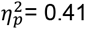), with higher values observed when preparing to reach targets in the direction of pectoralis shortening. For the posterior deltoid, there was also a main effect of preparatory delay (long>short; F_1,15_ = 6.6, p = 0.021, 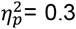), and an interaction effect between delay and target distance, with F_1,15_ = 5.2, p = 0.04 and 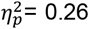. However, Tukey’s HSD test indicated that the impact of preparatory delay was present regardless whether the target was ‘near’ or ‘far’ (all p<0.031), but there were no significant differences involving target distance (i.e., short delay ‘far’ vs ‘near’: p=0.9; long delay ‘far’ vs ‘near’: p=0.094). Therefore, only in the pectoralis did target parameters affect pre-perturbation EMG activity. That is, there were no other main effects or interaction effects involving target parameters (or preparatory delay) in any of the analysed muscles (all p>0.056). It would therefore appear that, when preparing to reach with the non-dominant arm, target location impacts the pre-perturbation activity of the pectoralis in a statistically significant manner. Interestingly, target location did not affect pre-perturbation activity of the pectoralis of the dominant arm when performing the same (i.e., mirror-equivalent) delayed-reach task (Torell *et al.*, 2023). As described in the following sections, despite the presence of goal-directed differences in pre-perturbation EMG in the non-dominant limb, equivalent tuning of stretch reflexes in the nondominant limb was markedly reduced compared to that observed in the dominant limb.

#### The short-latency reflex epoch (SLR)

As mentioned in the Methods, it is well known that automatic gain-scaling due to muscle pre-loading limits the ability to unmask goal-directed modulation of SLR gains, due to the saturation of the goal-directed SLR response (Figures 2-4). Therefore, current analysis of the SLR responses focuses on the ‘no load’ and ‘unloaded’ conditions. That is, the ANOVA design in this case is 2(preparatory delay) x 2(load) x 2(target distance) x 2(target direction).

We found a consistent main effect of preparatory delay across all three analysed muscles (Fig. 5), with stronger SLR responses following a long delay (pectoralis: F_1,15_ = 13.1, p = 0.003, 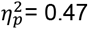; anterior deltoid: F_1,15_ = 11.0, p = 0.005, 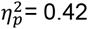; posterior deltoid: F_1,15_ = 12.1, p = 0.003, 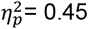). For the anterior deltoid, there were no other significant main or interaction effects on its SLR responses (all p>0.12). For the pectoralis and posterior deltoid, there was also a main effect of load on the SLR (pectoralis: F_1,15_ = 17.6, p = 0.0008, 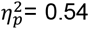; posterior deltoid: F_1,15_ = 12.4, p = 0.003, 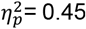), with weaker overall responses when the homonymous muscle was unloaded. For both these muscles, there was also a main effect of target direction, (pectoralis: F_1,15_ = 4.9, p = 0.043, 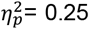; posterior deltoid: F_1,15_ = 12.1, p = 0.003, 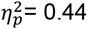), with stronger SLR responses when preparing to reach a target requiring shortening of the homonymous muscle (Fig. 5A and 5C, respectively). For the posterior deltoid alone, there was also a main effect of target distance (F_1,15_ = 10.6, p = 0.005, 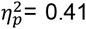), with significantly higher SLR responses when preparing to reach ‘far’ vs. ‘near’ targets, although the difference was quite small (difference between means = 0.0085 z).

**Fig. 5.**
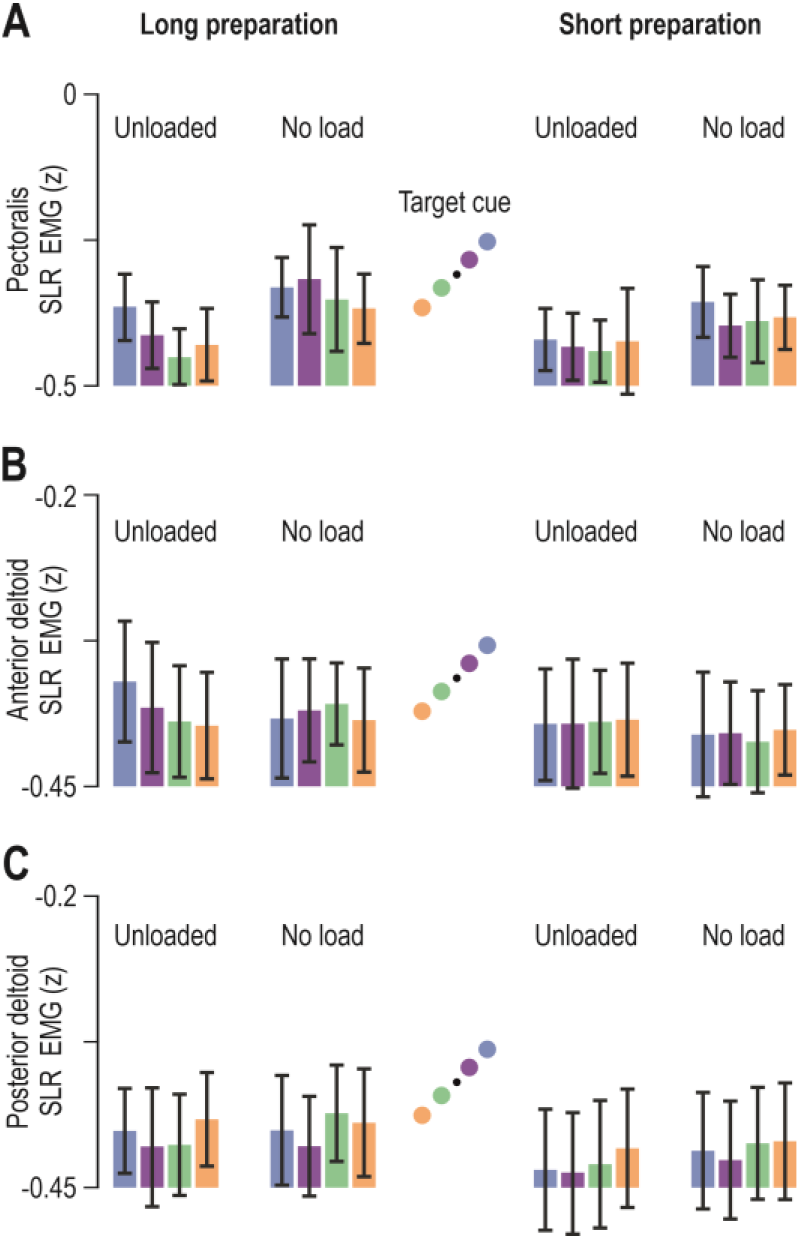
Lack of a consistent goal-directed modulation of SLR gains in the non-dominant limb. **(A)** The coloured bars represent mean pectoralis SLR EMG (z) across participants (N=16), and vertical lines represent 95% confidence intervals. Colour-coding as in previous figures. The axis range matches that used to illustrate the equivalent data for the dominant arm in Torell *et al.* (2023), for easier comparison. **(B)** As ‘A’ but representing the anterior deltoid muscle. There was no goal-directed modulation of anterior deltoid SLR. **(C)** As ‘A’ but representing the posterior deltoid.

To further describe the goal-directed modulation of the SLR, receiver operator characteristic (ROC) analyses were used. Since ANOVA did not show a consistent effect of target distance on SLR responses, the data was collapsed across target distance. Hence the ROC analyses concentrate on the impact of target direction. The relevant difference signals were created by contrasting the EMG curve observed when the cued targets were in the direction of homonymous muscle stretch vs. EMG curves of cued targets were in the direction of muscle shortening. In this way, the ROC curves were used to determine the time point at which the target direction related signals could be discriminated by an ideal observer. For the unloaded pectoralis (Fig. 6A), dog leg fits indicated deviance at 44 ms, for the no load condition at 46 ms and for the loaded condition at 53 ms. For the unloaded posterior deltoid (Fig. 6B), this occurred at 51 ms, for the no load condition at 44 ms and for the loaded condition at 54 ms. The onset times were also calculated for each participant individually (small red circles in Fig. 6A-B). The time point at which the ROC was above 0.75 was also identified and indicated as red vertical lines. For the unloaded pectoralis (Fig. 6A), this occurred at 63 ms, for the no load condition at 68 ms and for the loaded condition at 66 ms. For the unloaded posterior deltoid (Fig. 6B), this occurred at 69 ms, for the no load condition at 70 ms and for the loaded condition at 66 ms.

**Fig. 6.**
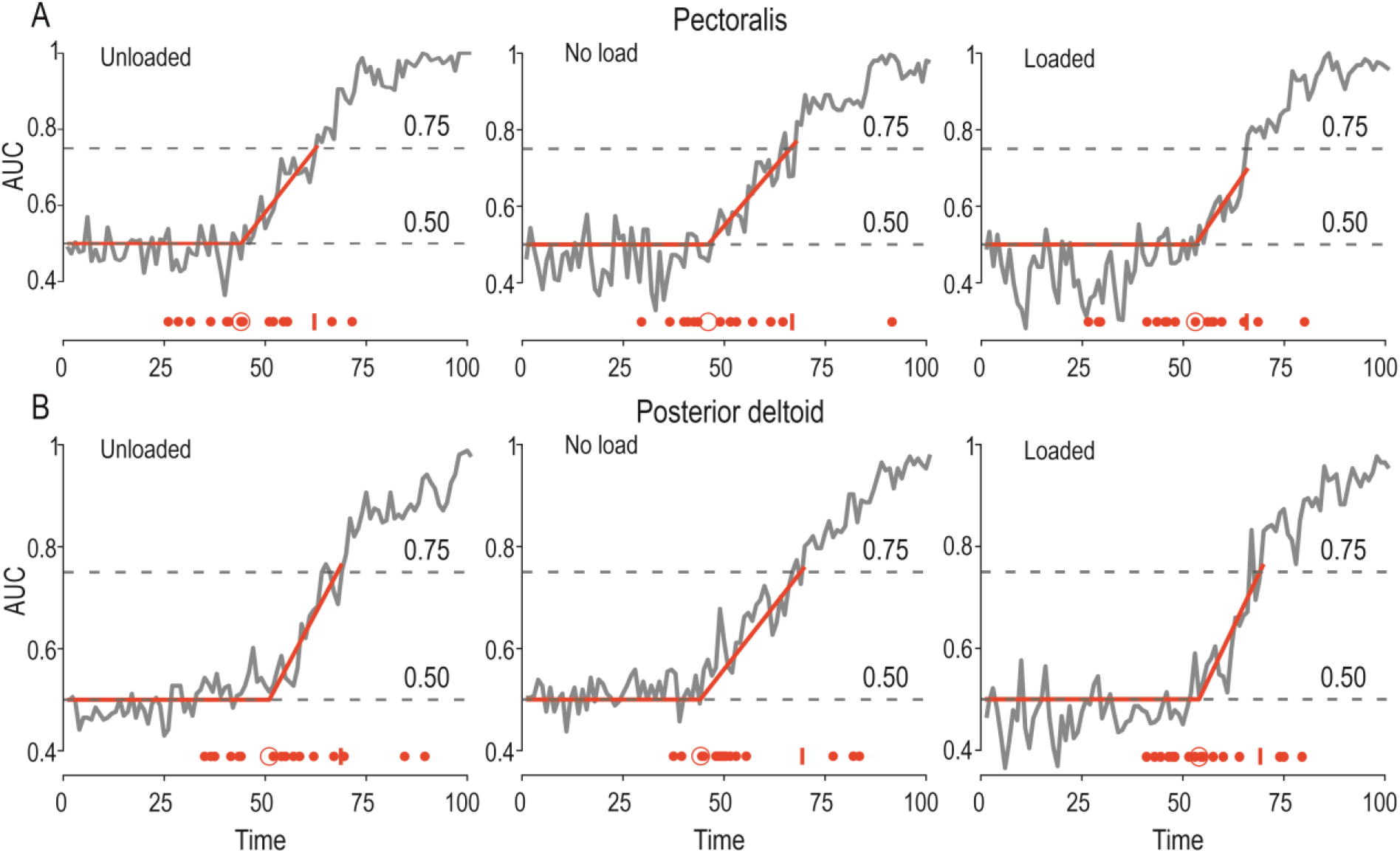
The time onset of SLR modulation. **(A)** The grey curve in each panel represents the area under the ROC, pertaining to pectoralis SLR modulation as a function of target direction, after experiencing one of the three load conditions (‘unloaded’, ‘no load’ and ‘loaded’) and a long preparatory delay (see Methods and Results for more details). Specifically, vertical axes represent the probability that an ideal observer could discriminate between the EMG difference curves. Each solid red line represents a dog leg fit which was applied to determine the onset of significant SLR modulation as a function of target direction (see also larger red circle at the bottom of each panel). The small red vertical line at the bottom of each panel represents the time point when the ROC area remained >0.75 for five consecutive time points (i.e., five consecutive ms). The smaller red dots represent the ROC result for each individual participant. **(B)** As ‘A’ but representing the posterior deltoid.

Compared to the previously performed ROCs in Torell *et al.* 2023 the modulation onset for unloaded pectoralis was 25 ms slower in the non-dominant arm. Similar figures for the no load condition 23 ms and similarly for the loaded pectoralis the modulation onset occurred 19 ms later in the non-dominant arm. For the posterior deltoid, the modulation onset for unloaded muscle occurred 23 ms slower. Similarly, for the no load condition 3 ms slower and for loaded posterior deltoid the onset occurred 4 ms later. In other words, the modulation differences were the strongest for the unloaded muscle.

The time point at which the ROC was above 0.75 were also compared. For the unloaded pectoralis the point of significant discrimination was delayed by 18 ms when the non-dominant arm was used. Similarly, for the no load condition significant discrimination could be achieved 8 ms faster using the dominant arm. For the unload pectoralis the difference was 9 ms, once again in favour of the dominant arm. For the unloaded posterior deltoid, significant discrimination could be achieved 17 ms faster for the dominant arm compared to the non-dominant arm. Similarly, for the no load condition and unloaded posterior deltoid was 9 ms and 3 ms, respectively. Once again, earliest goal-directed modulation in stretch reflex responses was evident for the for unloaded muscle.

#### The long-latency reflex epoch (LLR)

To examine LLR responses of muscles in the non-dominant limb, we used the complete ANOVA design of 2(preparatory delay) x 3(load) x 2(target distance) x 2(target direction). There was a main effect of load on the LLR responses of all three analysed muscles (Fig. 7; pectoralis: F_2,30_ = 12.7, p = 0.0001, 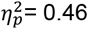; anterior deltoid: F_2,30_ = 14.7, p = 0.00004, 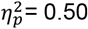; posterior deltoid: F_2,30_ = 22.2, p<10^-5^, 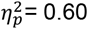). Tukey’s HSD test showed that LLR EMG activity was significantly higher when the pectorals was loaded (vs. unloaded: p=0.0004; vs. no load: p=0.0006), when the anterior deltoid was loaded (vs. unloaded: p=0.002; vs. no load: p=0.0002) and when the posterior deltoid was loaded (vs. unloaded: p=0.0001; vs. no load: p=0.0006). There was also a main effect of target direction on the LLR responses of all three muscles, with higher gains evident when preparing to reach targets associated with homonymous muscle shortening (pectoralis: F_1,15_ = 60.0, p<10^-5^, 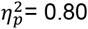; anterior deltoid: F_1,15_ = 22.6, p = 0.0003, 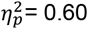; posterior deltoid: F_1,15_ = 74.0, p<10^-5^, 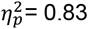). For the anterior deltoid, there was also a significant main effect of preparatory delay (F_1,15_ = 8.1, p = 0.012, 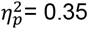), with stronger responses following a long preparatory delay. There was also a main effect of target distance on anterior deltoid LLR (F_1,15_ = 6.3, p = 0.024, 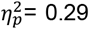), with stronger responses for ‘far’ vs. ‘near’ targets, but no such main effect was evident for the pectoralis or posterior deltoid (p=0.43 and p=0.13, respectively). In short, across all analysed muscles, there were consistent main effects of load and target direction on LLR responses.

**Fig. 7.**
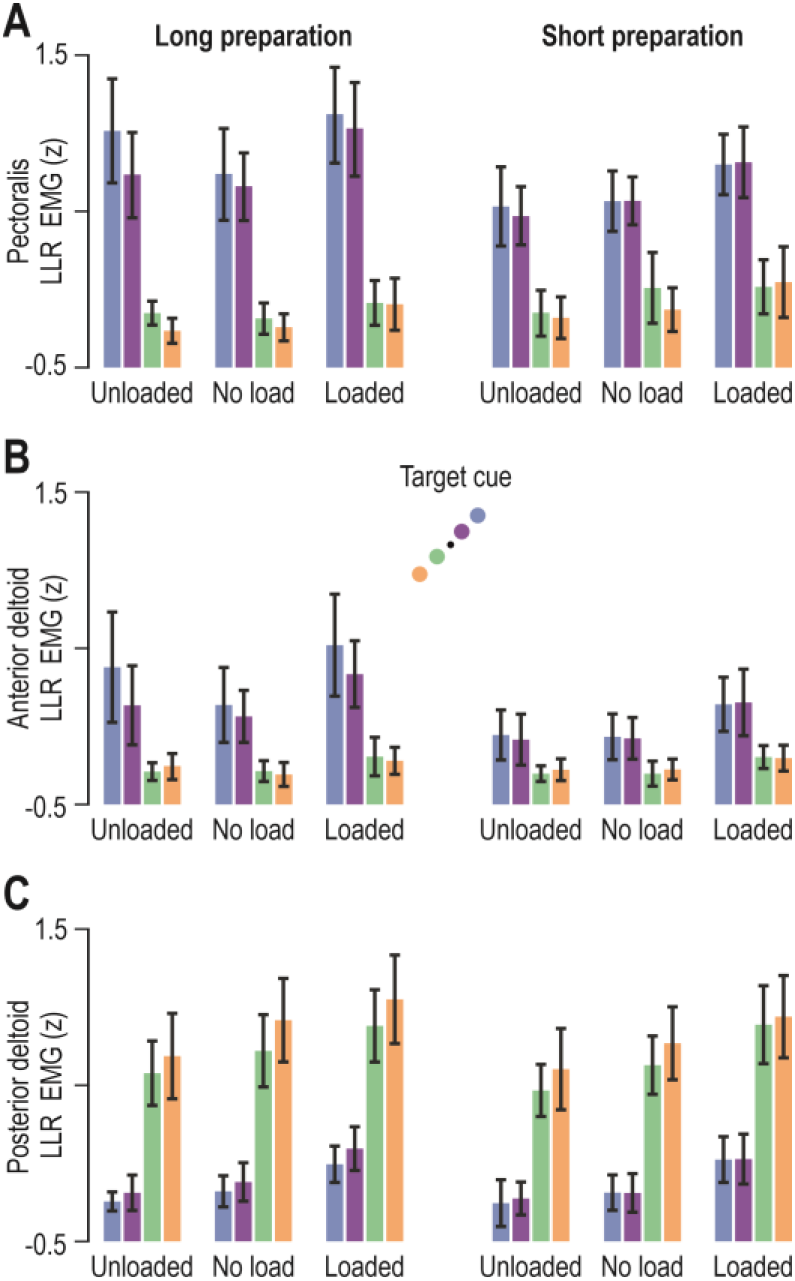
Goal-directed modulation of LLR gains in the non-dominant limb. **(A)** The coloured bars represent mean pectoralis LLR EMG (z) across participants (N=16), and vertical lines represent 95% confidence intervals. Colour-coding as in previous figures (see also schematic in ‘B’). ANOVA analyses indicated a significant impact of target direction on pectoralis LLR gains. **(B)** As ‘A’ but representing the anterior deltoid muscle. Similar LLR modulation patterns were observed as for the pectoralis. **(C)** As ‘A’ but representing the posterior deltoid. Equivalent LLR modulation patterns were observed as for the pectoralis.

In addition to the main effects, there were also significant two-factor interaction effects on the LLR responses. The ANOVA indicated a significant interaction effect between delay and target direction for both the pectoralis and anterior deltoid (pectoralis: F_1,15_ = 27.5, p = 0.00001, 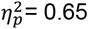; anterior deltoid: F_1,15_ = 20.7, p = 0.0004, 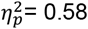). For both these muscles, Tukey’s HSD showed that the impact of preparatory delay on LLR responses (long > short) was evident only when preparing to reach targets requiring shortening of the homonymous muscle (i.e., blue and purple in Fig. 7A-B; p<0.008). In addition, there was a significant interaction effect between target distance and target direction on pectoralis and posterior deltoid LLR responses (pectoralis: F_1,15_ = 5.4, p = 0.035, 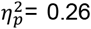; posterior deltoid: F_1,15_ = 21.2, p = 0.0003, 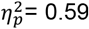). Tukey’s HSD test showed that, for the pectoralis, there was some impact of distance (‘far’ > ‘near’) only when comparing targets of a different direction (p>0.23 for same-direction comparisons). In addition, for the posterior deltoid, Tukey’s HSD indicated a significant impact of target distance on LLR responses (‘far’ > ‘near’) when preparing to reach targets associated with homonymous muscle shortening (p=0.001; orange vs. green in Fig. 7C).

For the pectoralis muscle, there was an additional interaction effect between preparatory delay and load (F_2,30_ = 4.0, p = 0.028, 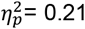) with Tukey’s HSD test showing that the impact of delay on pectoralis LLR (long > short) was evident when the pectoralis was unloaded (p=0.012). Last, there was an interaction effect between load and target direction on anterior deltoid LLR (F_2,30_ = 7.9, p = 0.002, 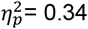). Tukey’s HSD showed that the majority of the two-factor interactions to be significant, except for comparisons of targets placed in the direction requiring muscle stretch (all p > 0.10). Nevertheless, despite the various interaction effects, it is rather clear that LLR gains in the non-dominant upper limb were primarily influenced by target direction, as can be appreciated by visually inspecting Figure 6.

#### Reflex tuning in the dominant vs. non-dominant upper limb

To compare goal-directed tuning of stretch reflex responses in the dominant vs. non-dominant arm, we use the data recorded for the purposes of this study and additional data published recently (Torell *et al.*, 2023), where a separate group of participants performed the equivalent experiment using their right dominant arm. Because target direction was the primary factor shaping reflex gains in both the dominant and non-dominant arm, the following analyses use data that were collapsed across target distances (i.e., green/orange were combined, as were blue/purple; e.g., Fig. 7), so that goal only represents target direction in this case. Moreover, in order to simplify analyses and produce a measure that is more directly representative of goal-dependent tuning, the responses observed when preparing to stretch the homonymous muscle (i.e., preparing to reach a target associated with muscle lengthening) were subtracted from the responses observed when preparing to shorten the homonymous muscle (see schematics in Figs. 8-9). As the anterior deltoid of the non-dominant arm showed no goal-directed modulation of SLR gains (Fig. 5B), the following analyses are limited to the posterior deltoid and pectoralis muscle.

**Fig. 8.**
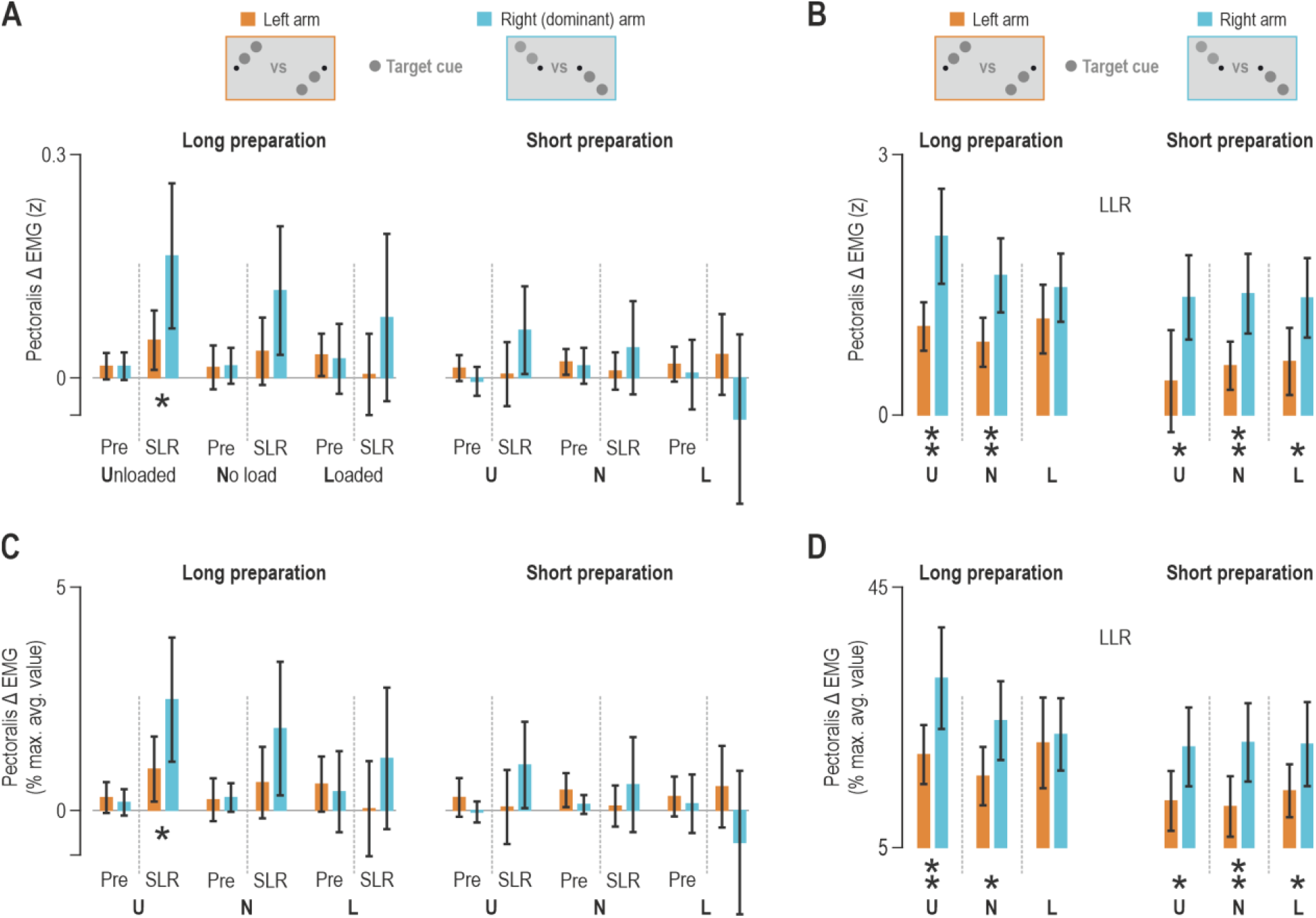
Goal-directed tuning of pectoralis stretch reflexes in the dominant vs. non-dominant limb. **(A)** The brown bars represent mean goal-directed change in pectoralis EMG activity (z) of the non-dominant arm, across participants (N=16). The relevant values for each individual muscle were generated by subtracting EMG activity observed in trials where participants prepared to stretch the homonymous muscle from the activity observed when preparing muscle shortening (i.e., activity in trials where reaching the cued target required shortening of the homonymous muscle; see also schematics). Cyan bars represent the corresponding EMG data of a group of participants (N=14) who performed the equivalent delayed-reach task using their dominant/right limb. Throughout, vertical lines represent 95% confidence intervals, ‘Pre’ represents the pre-perturbation epoch, single stars represent a significant difference following an independent t-test (brown vs. cyan) at alpha=0.05, and double stars indicate statistical significance at alpha=0.01. **(B)** As ‘A’ but representing the LLR response. **(C-D)** As ‘A-B’ but here EMG activity from each individual muscle was normalized as a proportion (%) of the maximal mean value across trials (see Methods for more details).

**Fig. 9.**
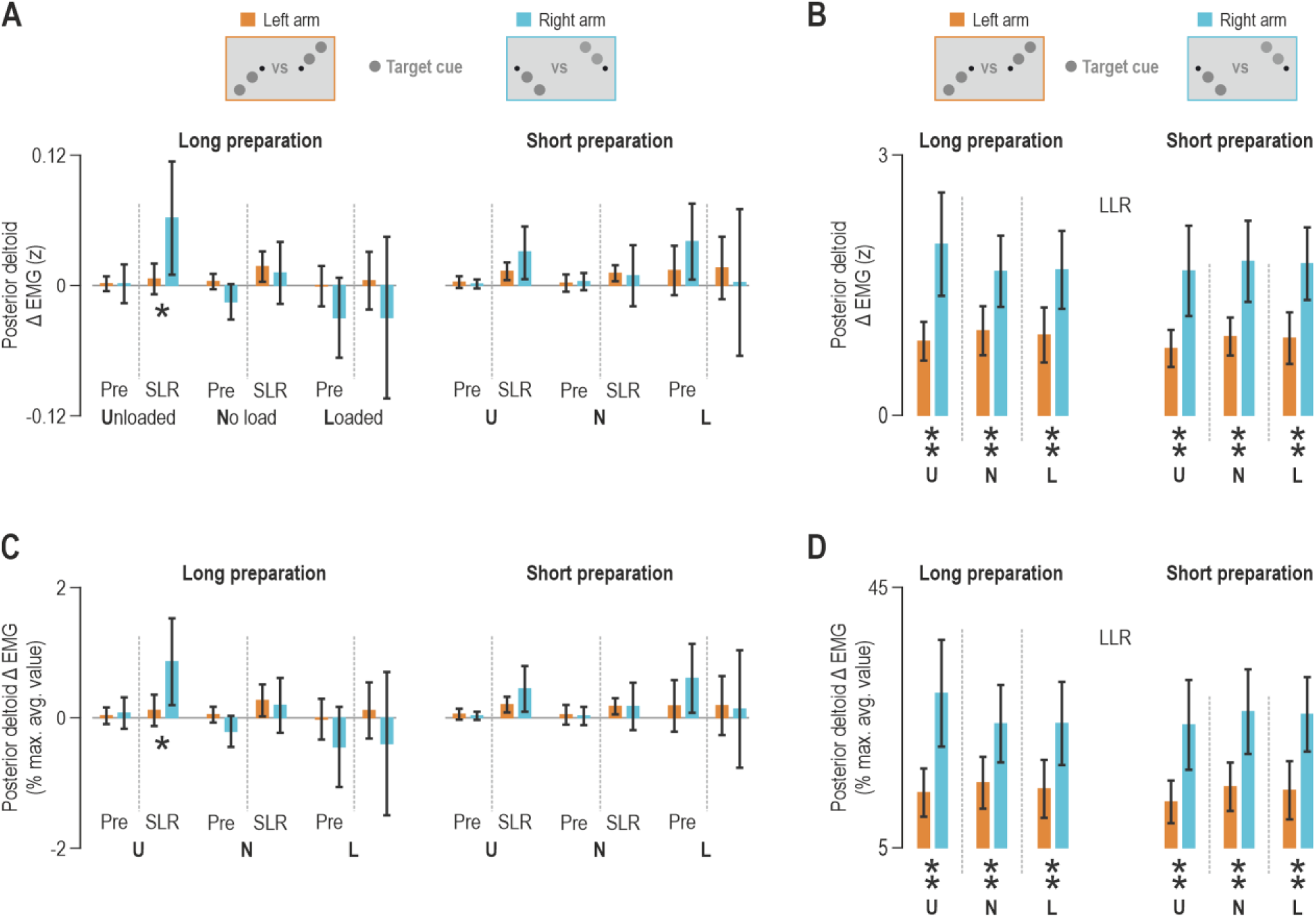
Goal-directed tuning of posterior deltoid stretch reflexes in the dominant vs. non-dominant limb. **(A)** The brown bars represent mean goal-directed change in posterior deltoid EMG activity (z) of the non-dominant arm, across participants (N=16). The relevant values were generated as per Figure 8A-B. (see also schematics). Cyan bars represent the corresponding EMG data of a separate group of participants (N=14) who performed the equivalent delayed-reach task using their dominant/right limb. Throughout, vertical lines represent 95% confidence intervals, ‘Pre’ represents the pre-perturbation epoch, single stars represent a significant difference following an independent t-test (brown vs. cyan) at alpha=0.05, and double stars indicate statistical significance at alpha=0.01. **(B)** As ‘A’ but representing the LLR response. **(C-D)** As ‘A-B’ but here EMG activity from each individual muscle was normalized as a proportion (%) of the maximal mean value across trials.

With regard to the unloaded pectoralis major, an independent t-test with z-normalized data indicated stronger goal-directed tuning of SLR gains in the dominant arm following a long preparatory delay (t_28_=2.4, p=0.022), with no corresponding difference in pre-perturbation activity (t_28_=0.0009, p=0.99; Fig. 8A, leftmost bars). The difference in pectoralis SLR between the dominant and non-dominant limbs failed to reach significance in the no-load condition following a long delay (t_28_=1.87, p=0.07). However, a single-sample t-test showed that pectoralis SLR responses from the dominant limb (cyan) were significantly different from zero (t_13_=2.93, p=0.012), confirming the presence of a goal-directed tuning in this case, whereas this was not so for SLR responses from the non-dominant limb (t_15_=1.68, p=0.1). There were no differences in SLR tuning when the pectoralis muscle was loaded (t_28_=1.37, p=0.18), nor under any load condition when the preparatory delay was short (right panel in Fig.8A; brown vs. cyan unloaded: p=0.09; no load: p=0.3; loaded: p=0.13). Note the gradual decrease in SLR gain of the dominant pectoralis from left-to-right in Figure 8A (cyan bars), highlighting earlier findings that longer preparation and unloading (i.e., ‘assistive’ loading) promote the goal-directed modulation of stretch reflex gains.

Figure 8B displays the goal-directed LLR responses of the pectoralis in the dominant and non-dominant limb. When the preparatory delay was long, there was significantly stronger goal-directed tuning of LLR gains in the dominant arm when the muscle was unloaded (t_28_=3.78, p=0.0008) and when there was no load (t_28_=3.31, p=0.003). Interestingly, there was no corresponding difference in LLR tuning when the pectoralis of the dominant and non-dominant arms was loaded (t_28_=1.38, p=0.18). In contrast, when the preparatory delay was short, pectoralis LLR gains were significantly higher in the dominant arm under all load conditions (unloaded: t_28_=2.66, p=0.013; no load: t_28_=3.43, p=0.002; loaded: t_28_=2.64, p=0.013).

Equivalent significant differences were obtained for the SLR and LLR when the %-normalized pectoralis activity was analysed (Fig. 8C-D). Specifically, for the unloaded pectoralis, a t-test indicated stronger goal-directed tuning of SLR gains in the dominant arm following a long preparatory delay (t_28_=2.2, p=0.035), with no corresponding difference in pre-perturbation activity (t_28_=-0.5, p=0.62; Fig. 8C, leftmost bars). When the preparatory delay was long, there was significantly stronger goal-directed tuning of LLR gains in the dominant arm when the muscle was unloaded (t_28_=2.89, p=0.007) and when there was no load (t_28_=2.47, p=0.02). When the preparatory delay was short, pectoralis LLR gains were significantly higher in the dominant arm under all load conditions (unloaded: t_28_=2.37, p=0.025; no load: t_28_=2.81, p=0.009; loaded: t_28_=2.07, p=0.047).

Similar reflex tuning patterns are evident for the posterior deltoid (Fig. 9). With regards to the unloaded muscle, we found stronger goal-directed tuning of SLR gains in the dominant arm following a long preparatory delay (t_28_=2.37, p=0.025), again with no corresponding difference in pre-perturbation activity (t_28_=-0.017, p=0.97; Fig. 9A, leftmost bars). There was no difference in posterior deltoid tuning between the dominant and non-dominant limb in the no-load condition (t_28_=-0.41, p=0.67) or loaded condition (t_28_=-0.98, p=0.37), following a long delay. When the preparatory delay was short, there were no differences in SLR tuning under any load condition (right panel in Fig.9A; unloaded: p=0.14; no load: p=0.86; loaded: p=0.68). Figure 9B displays the goal-directed LLR responses of the posterior deltoid. Regardless of preparatory delay duration or load condition, goal-directed tuning of posterior deltoid LLR was stronger in the dominant arm (from left to right in Figure 8D: t_28_=3.99, p=0.0004; t_28_=3.02, p=0.005; t_28_=2.98, p=0.006; t_28_=3.58, p=0.0013; t_28_=3.76, p=0.0008; t_28_=3.66, p=0.001).

Equivalent significant differences were obtained for the SLR and LLR when the %-normalized posterior activity was analysed (Fig. 9C-D). Specifically, for the unloaded posterior deltoid, there was stronger goal-directed tuning of SLR gains in the dominant arm following a long preparatory delay (t_28_=2.4, p=0.024), and no corresponding difference in pre-perturbation activity (t_28_=0.034, p=0.74; Fig. 9C, leftmost bars). Regardless of preparatory delay duration or load condition, goal-directed tuning of posterior deltoid LLR was stronger in the dominant arm (from left to right in Figure 9D: t_28_=3.82, p=0.0007; t_28_=2.79, p=0.009; t_28_=2.8, p=0.008; t_28_=3.48, p=0.0017; t_28_=3.42, p=0.0019; t_28_=3.5, p=0.0016). Overall, therefore, it appears that goal-directed tuning of stretch reflexes is stronger in the dominant upper limb, and this augmentation is particularly robust in the LLR epoch.

## Discussion

In this study we investigate the neural mechanisms that underlie the preparation for goal-directed movements in the non-dominant arm. The stretch reflexes in the non-dominant arm were examined using the same design used for the dominant arm (Torell *et al.*, 2023). Participants were presented with four targets varying in both direction and distance, and three background loads. After target presentation, a rapid perturbation was applied to the arm at one of two delays (short or long) to probe the preparatory tuning of stretch reflexes. We found small goal-directed differences in the background (pre-perturbation) muscle activity, but only minor differences in the short latency stretch reflex as a function of target location. The only consistent effect at the short latency was that of preparatory time, where we found stronger responses for the longer delay times. At the long latency interval, we found consistent effects of both load and target direction on the stretch reflex responses. Finally, we compared the goal-directed stretch reflex tuning found in the non-dominant arm with that previously seen in the dominant arm. The results clearly show much stronger goal-directed modulation of stretch reflex responses in the dominant arm, both in the short and long latency reflex intervals (‘SLR’ and ‘LLR’, respectively).

One possible explanation of the SLR and LLR amplitude differences could be differences in the alpha-motor neuron density projecting to the dominant and non-dominant arm. However, the amount of spinal motor neurons does not differ significantly between arms (Li *et al.*, 2015). In addition, intramuscular EMG recordings have shown that right-handed individuals have no dominance-related differences in single motor unit action potentials (Nelson *et al.*, 2003). On the other hand, genetics or long-term preferential use of the dominant arm seem to affect the muscle fibre composition. The proportion of Type I muscle fibres is significantly higher in the dominant arm of right-handed individuals (Fugl-Meyer *et al.*, 1982). Studies using surface EMG have also indicated asymmetric muscle composition related to dominance (Merletti *et al.*, 1994; Williams *et al.*, 2002). We propose that it is instead subtle differences in strategies of feedforward control between the dominant and non-dominant arm, particularly the differential control of gamma motor neurons, that gave rise to the observed asymmetries in SLR and LLR response amplitude. One potential limitation of the current study is that a different group of participants was examined compared to our previous study that documented reflex tuning in the dominant upper limb (Torell *et al.*, 2023). Although comparing the stretch reflex modulation in the left and right limbs of the same participants would likely provide stronger statistical power, the SLR and LLR differences between the dominant and non-dominant limbs in our study are nevertheless clear across all investigated muscles. Moreover, relevant analyses were conducted using normalized data, including normalization based on the maximal activation values observed in each individual muscle (Figs. 8-9).

Although there was evidence for a goal-directed modulation of reflex responses in the non-dominant arm, this tuning was clearly weaker than that observed in the dominant arm. However, we again found that the goal-directed modulation, particularly of the LLR, was more pronounced when sufficient preparatory time was allowed. Despite the similarity in the response patterns across both dominant and non-dominant limbs, a comparison shows much greater goal-directed tuning of the reflex gains in the dominant arm prior to reaching. This effect is particularly clear in the LLR epoch (Figs. 8B, 8D, 9B, 9D) where virtually every comparison showed stronger goal directed responses in the dominant arm. However, we also observed stronger goal directed tuning of SLR gains following a longer preparatory delay (Figs. 8A, 8C, 9A, 9C). The effect of preparatory delay on the goal-directed tuning of stretch reflex gains suggests that motor planning is not only composed of setting up the appropriate descending drive to the extrafusal muscles through alpha motor neuron control, but also tuning the gamma drive independently to provide goal-appropriate feedback. Since goal- and delay-directed modulation is evident for both SLR and LLR gains, this suggests a critical role for gamma motor neurons in movement preparation. Such changes in feedback gains take time to develop even for simple reaching movements, where a 250 ms delay shows much less modulation than 750 ms. A similar temporal evolution of LLR gains according to motor planning was shown in the random dot paradigm, where stronger motion coherence produced responses more appropriate for the subsequent movement (Selen, 2012).

The difference in goal-directed modulation of reflex responses between the dominant arm and non-dominant arm results was largest in the unloaded and no-load conditions in the long delay condition (although similar effects are visible in the loaded conditions, Fig. 8A, 8C). We have suggested that the strongest goal-directed modulation occurs when the muscle is unloaded because antagonist loading is accompanied by top-down reciprocal inhibition of the lower motor neurons of the muscle, including gamma motor neurons (Dimitriou, 2014). We have hypothesized that the stronger goal-directed modulation develops as the independent goal-directed control of dynamic gamma motor neurons occurs on top of this blanket reciprocal inhibition of lower motor neurons that accompanies muscle unloading (Torell *et al.*, 2023). More specifically we suggest that preparatory activity before a planned voluntary movement is involved in setting this goal-dependent tuning of muscle spindles just prior to a movement (Papaioannou & Dimitriou, 2021). This effect is dampened by directly loading the muscle as gain-scaling dominates the overall SLR responses, limiting the ability to detect goal-directed tuning (Torell *et al.*, 2023). This is different to the historical view that preparatory activity represents a sub-threshold version of movement-related cortical activity (Tanji & Evarts, 1976). With such a hypothesis, we might be concerned that the no-load or even the unloaded conditions might exhibit different sub-threshold activity of the motor neuron pools that could affect the reflex gains (Beddingham & Tatton, 1984; Matthews, 1996; Pruszynski *et al.*, 2009). However, recent work has contradicted the idea of sub-threshold preparation, showing instead that preparatory activity sets an initial dynamical state that promotes execution of the planned movement (Churchland *et al.*, 2010). While preparation lowers reaction time (Rosenbaum, 1980) and benefits movement quality (Ghez *et al.*, 1997), it has been shown that preparation is mechanistically independent from movement initiation (Haith *et al.*, 2016). Such studies motivated the possibility that preparatory activity might reflect the control of proprioceptive sensory feedback, by setting the appropriate goal-directed modulation of the fusimotor system. This possibility was directly tested using microneurography, showing that preparation for moving to a visual target sets changes in spindle stretch sensitivity, but that this affect takes time to fully develop (Papaioannou & Dimitriou, 2021). To assess whether this difference in spindle activity could affect stretch reflexes, goal-directed modulation was confirmed in a delayed reaching task (Papaioannou & Dimitriou, 2021). Here we use an equivalent experimental design to investigate differences in the goal-directed modulation of stretch reflexes across the dominant and non-dominant arms. While we cannot directly confirm these differences arise due to laterality of fusimotor control across the two arms, this does provide a strong prediction that can be directly tested in future studies.

The dominant arm exhibited stronger goal-directed modulation of stretch reflexes, suggesting that the dominant arm is better at anticipating and controlling reflex muscle stiffness according to the movement goal. Therefore, the dominance differences seen during voluntary movement, e.g., faster reaction time and better fine motor-skills (Annett *et al.*, 1979; Shen & Franz, 2005), are also visible in feedback control. Previous research has suggested that the non-dominant side may rely more heavily on proprioceptive feedback (Goble & Brown, 2008b). This fits with the theory that the two hands play different roles in motor control (Sainburg, 2005; Sainburg, 2014; Sainburg & Kalakanis, 2000; Bagesteiro & Sainburg, 2003), with the non-dominant hand controlling position and the dominant hand controlling trajectory. The current theory of motor lateralization is heavily based on findings in individuals with unilateral brain damage (Schaefer *et al.*, 2009, 2012). The theory suggests that right-handed individuals have a dominant (left) hemisphere that accounts for predictive, feedforward control and movement planning, while the non-dominant hemisphere accounts for feedback-mediated error correction. This in turn implies that the non-dominant hand is under impedance control while the dominant hand is under trajectory control through internal predictive models.

The CNS has the capacity to selectively control muscle spindle sensitivity, an ability which has also been shown in completely relaxed subjects (Ribot-Ciscar *et al*., 2000). In this way, both hemispheres may contribute significantly to the control of goal-directed movements, where the ipsilateral hemisphere is inhibited using a signalling pathway mediated via the spinal cord (Chen *et al.*, 1997; Fisk & Goodale, 1988; Haaland & Harrington, 1989; Winstein & Pohl, 1995). Transcranial magnetic stimulation has shown that dominance is associated with asymmetrical excitability of the corticospinal system (De Gennaro *et al.*, 2004). De Gennaro *et al.* found that the dominant hemisphere had lower thresholds in left-handed participants, and larger amplitudes of motor evoked potential (MEP) in right-handed participants. Conflicting findings have been presented where MEP was found to be unrelated to arm dominance and instead dependent on the length of the upper extremity (Livingston *et al.*, 2010).

Here we find differences in the laterality of goal-directed modulation of stretch reflexes in our delayed-reach task. Specifically, we find higher modulation of both SLR and LLR gains according to the future movement goal in the dominant arm. In contrast, there have been limited differences in laterality seen in postural tasks (Walker & Perreault, 2015; Maurus *et al.*, 2021). Walker and Perreault (2015) found little evidence for modulation in reflex sensitivity according to arm dominance across the whole population across the age groups. In a similar postural study, Maurus and colleagues (2021) examined the reflex sensitivity and directionality between the dominant and non-dominant limbs. They found no evidence for differences in the pattern of reflexes between the two limbs, and little difference in the magnitude of the responses. Although this study (Maurus *et al.*, 2021) predicted that the non-dominant limb would exhibit stretch reflexes aligned with single joint motion due to a possible reliance on impedance control, it has long been known that the control of limb impedance requires impedance to be coordinated across all joints to maintain system stability (McIntyre *et al.*, 1995; Franklin & Milner 2003; Franklin *et al.*, 2007), and therefore likely also requires accurate internal models of limb dynamics (Franklin *et al.*, 2008; Tee *et al.*, 2010). We therefore argue that a difference between impedance control and internal model control would be unlikely to show directionality differences in this postural task (Maurus *et al.*, 2021).

Nevertheless, both previous studies showed limited differences in the laterality of stretch reflexes, in contrast to our results, which likely reflect the major differences in the task design. First, both postural task studies pre-load the limbs and muscles limiting the modulation of stretch reflexes (Dimitriou, 2018; Torell *et al.*, 2023). Secondly, these postural tasks simply require postural maintenance regardless of the nature of the perturbation, whereas our task required participants to plan a movement towards one of multiple targets. Such movement planning appears to activate differential control of the gamma motor neurons to tune muscle spindle and reflex feedback to the upcoming movement (Papaioannou & Dimitriou, 2021). Such goal-directed modulation of stretch reflexes may reflect the development and tuning of feedforward predictive models which Sainburg and colleagues proposed to be the case for the dominant limb (Sainburg, 2005; Sainburg, 2014; Sainburg & Kalakanis, 2000; Bagesteiro & Sainburg, 2003).

Goal-directed tuning of SLR gain is likely the result of changes in gamma drive. The old view that muscle spindle sensitivity is mainly associated with α-γ-coactivation (Vallbo, 1970; Vallbo *et al.*, 1979) is moving toward a more dynamic view where the fusimotor system allows advanced signal-processing to occur at the periphery (Dimitriou, 2022). The observed effects of target direction and arm dominance on the SLR (and LLR) suggest that the dominant arm displays better independent fusimotor control. At the SLR epoch, we have found stronger goal-directed modulation of responses when the muscle is unloaded compared even to when there is no load applied to the limb. While such effects are most prominent in the dominant limb (Torell *et al.*, 2023), we also find similar results here in the non-dominant limb following a longer preparatory delay. We hypothesize that this arises as the loading of the antagonists is accompanied by top-down reciprocal inhibition of lower motor neurons of the muscle, including gamma motor neurons (Dimitriou, 2014). This likely contributes to stronger goal-directed effects as the independent goal-directed control of dynamic gamma motor neurons occurs on top of this blanket reciprocal inhibition of lower motor neurons. Indeed, there is evidence that such goal-directed tuning of muscle spindles may occur primarily through changes in dynamic gamma drive to primary muscle spindles, as there is no evidence for preparatory goal-directed tuning of secondary muscle spindles (Papaioannou & Dimitriou, 2021).

Overall, the current study supports previous findings that sufficient preparation before a planned movement and muscle unloading (i.e., antagonist loading) facilitates goal-directed modulation of stretch reflexes, in both the short and long latency intervals. (Papaioannou & Dimitriou, 2021; Torell *et al.*, 2023). Here, we extend the work by showing goal-directed tuning of stretch of reflexes is significantly reduced in the non-dominant limb compared to the dominant limb. Such differences may reflect the different control involved depending on limb laterality and highlights a potential role for independent gamma drive in the higher skill and function of the dominant limb.

## Notes

### Competing Interest Statement

The authors have declared no competing interest.

### Summary of Updates

This version contains: * Significance statement has changed to Key points * Updated the introduction with additional references. * New normalization technique and comparison between normalization techniques * Fig. 7 revised and now called Fig. 8 and Fig. 9. * ROC-curves to compare reflex modulation onset between the dominant and non-dominant upper limb. * Fig. 7 now contains the ROC-curves. * Extended discussion of our results.

